# Genome Scans for Selection and Introgression based on *k*-nearest Neighbor Techniques

**DOI:** 10.1101/752758

**Authors:** Bastian Pfeifer, Nikolaos Alachiotis, Pavlos Pavlidis, Michael G. Schimek

## Abstract

In recent years, genome-scan methods have been extensively used to detect local signatures of selection and introgression. Here, we introduce a series of versatile genome-scan methods that are based on non-parametric *k*-nearest neighbors (kNN) techniques, while incorporating pairwise Fixation Index (*F*_*ST*_) estimates and pairwise nucleotide differences (*d*_*xy*_) as features. Simulations were performed for both positive directional selection and introgression, with varying parameters, such as recombination rates, population background histories, the proportion of introgression, and the time of gene flow. We find that kNN-based methods perform remarkably well while yielding stable results almost over the entire range of *k*. We provide a GitHub repository (*pievos101/kNN-Genome-Scans*) containing R source code to demonstrate how to apply the proposed methods to real-world genomic data using the population genomics R-package PopGenome.

## Introduction

The last years have seen great advances in whole genome sequencing and population genomics methods to detect DNA fragments affected by natural selection. Genomic regions under selection are assumed to be rare and thus can be considered as anomalies which deviate from the overall population structure. These anomalies are of great interest as they may act as a major force in the adaptation of populations to their environments during evolution. One of the most widely applied statistics to detect such regions is the Fixation Index (*F*_*ST*_), which was originally proposed as a measure of population differentiation under the Wright-Fisher model (Wright, 1949).

Several variations of the *F*_*ST*_ are used by the population genomics community (Hudson *et al.*, 1992; Weir, 1996; Weir and Cockerham, 1984). In general, a high *F*_*ST*_ can be an indication of positive directional selection. However, in cases where the neutral population background history (the neutral distribution of *F*_*ST*_) is not known, hypothesis testing of neutral evolution is nearly impossible. This especially applies in cases where the population history deviates from the Wright-Fisher model, or when hierarchical structure is introduced to the system. In such cases, results based on *F*_*ST*_ are no longer reliable (Excoffier *et al.*, 2009; Foll and Gaggiotti, 2008).

Also, the *F*_*ST*_ estimate was introduced as a model parameter in Bayesian approaches, and inferred via computationally intensive Markov-Chain-Monte-Carlo (MCMC) simulations. In such approaches, a common migrant pool is modeled as a Dirichlet distribution, and the genome-wide neutral signal is captured in a logistic regression model with a specific parameter shared by all populations. One of the most prominent methods is implemented in the BayeScan software (Foll and Gaggiotti, 2008), which is built upon the works of Beaumont and Nichols (1996) and Beaumont and Balding (2004). It has been reported, however, that these methods suffer from a high False Discovery Rate (FDR) (De Villemereuil and Gaggiotti, 2015; Duforet-Frebourg *et al.*, 2014, 2015). More recently, published approaches based on Principal Component Analyses (PCA) address some of these shortcomings (Duforet-Frebourg *et al.*, 2015; Luu *et al.*, 2017), and at the same time they are computationally less demanding.

Another topic of great interest is the investigation of hybridization and the detection of the related introgressed regions in whole genome scans. Hybridization between species is increasingly recognized as an evolutionary force in which species share genetic information across the species boundary. There has been an explosion of available methods in this area in recent years. Currently, the most widely applied methods are the ABBA-BABA family of methods which are based on a four-taxon system (Durand *et al.*, 2011; Green *et al.*, 2010; Martin *et al.*, 2014; Pfeifer and Kapan, 2019), where the fourth taxon acts as the outgroup. Since an outgroup is not always available, several other approaches based on fewer taxa were also introduced (Geneva *et al.*, 2015; Hahn and Hibbins, 2019; Hibbins and Hahn, 2019).

We believe that the ability of techniques based on *k*-nearest neighbors (kNN) to detect local signatures of selection or introgression is widely underestimated, and detailed investigations on the use of these techniques are yet to be reported. The kNN-based approaches are among the oldest unsupervised machine learning techniques and have been widely applied in almost all areas of data-driven research. In this paper, we make use of kNN-based techniques while incorporating pairwise *F*_*ST*_ estimates as features. We study the ability of these approaches to detect local signatures of directional selection and introgression under a comprehensive set of simulation scenarios. In the case of introgression, we also investigate the usage of pairwise nucleotide differences (*d*_*xy*_) as features because it has been reported that *F*_*ST*_ estimates may lead to false positives when diversities within populations are low (Cruickshank and Hahn, 2014).

We compare the accuracy of these approaches with recently published genome-scan methods, and finally showcase the use of the kNN approaches to detect positively selected regions in the human genomics data made available by the 1000 genomes project (1000 Genomes Project Consortium and others, 2015).

## New Approaches

### kNN Techniques using *F*_*ST*_ as Features

The key idea of the kNN approach (Ramaswamy *et al.*, 2000) is to calculate the distances between a given data point and its *k* nearest neighbors. Data points at high distance from their neighborhood are considered as outliers. In this work, we use pairwise *F*_*ST*_ estimates, as proposed by Hudson *et al.* (1992) and recommended by Bhatia *et al.* (2013), and incorporate them as features into kNN-based algorithms. Consequently, the population pairwise *F*_*ST*_ estimates define a genomic region as a data point embedded into an *m*-dimensional numerical space, where *m* is the total number of possible population pairwise comparisons (*m*= *n*_*p*_(*n_p_ −*1)/2), and *n*_*p*_ is the total number of populations analyzed. Thus, the population structure of each genomic region is represented by an *F*_*ST*_ vector of length *m*. The kNN outlier score for a given genomic region *x* is calculated as follows:

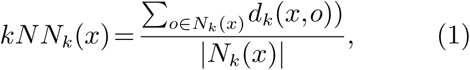

where *N*_*k*_(*x*) is the *k*-nearest neighbor set of the genomic region *x*, and *d*_*k*_(*x,o*) defines the dissimilarity between the genomic regions *x* and *o*. It is calculated as the euclidean distance between the pairwise *F*_*ST*_ vectors 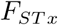 and 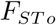:

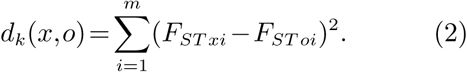

The basic kNN approach was slightly modified by the weighted-kNN approach (Angiulli and Pizzuti, 2002, 2005), which takes into account the overall distance from a data point to its neighborhood by calculating the sum of distances instead of the arithmetic mean. Another way of calculating the distances is implemented in ODIN (Outlier Detection using Indegree Number) (Hautamaki *et al.*, 2004), which infers outliers based on a kNN graph.

A wide range of methods have been developed to also account for local outlierness. The best known one is LOF (Local Outlier Factor) (Breunig *et al.*, 2000), which is based on the concept of *local reachability density* (*lrd*) of the *k*-nearest neighbors. In this context, a data point is considered to be an outlier when its density is much smaller than the densities of its neighbors. The *lrd* is defined as

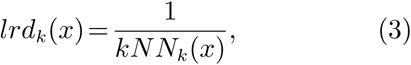

and the LOF can be calculated as

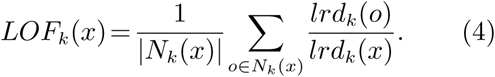

The LOF algorithm and the corresponding *lrd* concept was later modified in several ways. For example, the simplified-LOF (Schubert *et al.*, 2014) uses the basic kNN distances instead of the LOFs reachability distance. COF (Connectivity-based Outlier Factor) (Tang *et al.*, 2002) modifies the density estimation of the simplified-LOF to account for the connectedness of a neighborhood via a minimum spanning tree (MST). Another tool is called LoOP (Local Outlier Probabilities) (Kriegel *et al.*, 2009), which adopts normalized local density scores based on the quadratic mean. Therefore, the scores are strictly within the [0,1] interval, and can be interpreted as *p*-values. LDOF (Local Distance-based Outlier Factor) (Zhang *et al.*, 2009) uses the relative distance from a data point to its neighbours, measuring how many data points deviate from their scattered neighbourhood. The ABOD approach addresses the so-called *curse of dimensionality* problem by comparing the angles between pairs of distance vectors. FastABOD (Fast Angle-Based Outlier Detection) is a faster variant of ABOD (Kriegel *et al.*, 2008). LDF (Local Density Factor) (Latecki *et al.*, 2007) replaces LOFs density estimation by a variable-width Gaussian kernel density estimation (KDE). INFLO (Influenced Outlierness) (Jin *et al.*, 2006) takes into account also the reverse nearest neighborhood set when calculating the local density scores.

In fact, there exist many more of these type of anomaly detection algorithms which we have not mentioned here. In this work, we focus on a subset of kNN approaches as suggested by Campos *et al.* (2016).

### The Selection of *k*

In classification problems, the parameter *k* can be inferred, for instance, by cross-validation. However, it is well known that the inference of an appropriate *k* in a purely unsupervised setting is a challenging task, and highly depends on the data analyzed. This challenge especially arises in studies where the goal is the detection of local outliers. However, here we use the kNN-based methods for global outlier detection, with the aim to distinguish between signals of neutral evolving genomic regions and outlier regions subject to selection or introgression. Thus, the choice of *k* may not have a big influence on the outcomes as long as the value is not too small.

There are two main requirements that an adequate *k* should fulfill. First, the outlier scores of the corresponding kNN methods need to be reasonable, which we think is fulfilled when the ranks of the kNN scores approximately align with the findings of well established methods. Second, the *k* should be settled in a stable *k* region. We refer to a stable *k* region when the ranks of the corresponding kNN scores within that region are highly correlated. To address the former requirement we simply use pairwise *F*_*ST*_ estimates as features for the kNN-based methods. Pairwise *F*_*ST*_ is successfully applied in genomic scans to detect positive selection as well as introgression. In this work, we use the *F*_*ST*_ estimate as proposed by Hudson *et al.* (1992) and recommended by Bhatia *et al.* (2013). To address the latter requirement we propose the following approach:

First, calculate the kNN-scores (**s**_*i*_) for *n*_*k*_ =100 sequentially sampled *k*s from [2*,n_r_ −*1], where *n*_*r*_ is total number of genomic regions:

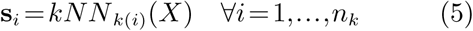

Second, calculate Kendalls tau (*τ*) correlation coefficients

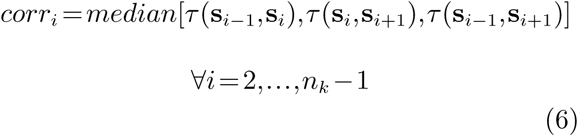

where

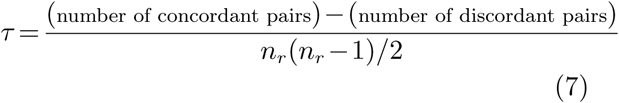

Third, from the correlation vector *corr* infer the longest connected *k* region with *corr >* 0.90, and define the median of that region as the optimal *k*.

### Inferring the Type of Selection

The kNN-based methods based on pairwise *F*_*ST*_ features are reporting on anomalies as strong deviations from the overall population structure. Once the outlier are detected from the kNN-based outlier scores we suggest to remove the corresponding pairwise *F*_*ST*_ vectors and to calculate the *medoid* based on the remaining data points. We argue that the *medoid* is the most informative data point and sufficiently reflects the overall population structure. Subtracting the *medoid* from the outlier pairwise *F*_*ST*_ vectors will indicate which population or population pairs are affected by selection. We are pointing to these resulting vectors as the ∆*F*_*ST*_ selection effects. Positive entries of ∆*F*_*ST*_ are an indication for positive directional selection, whereas negative values point to introgression (reduced diversity due to gene-flow) or other types of selection which significantly reduce the diversity between populations, such as balancing selection.

## Results

### On the Power to Detect Positive Directional Selection

Simulations under positive directional selection (see ‘Material and Methods’) indicate that the kNN-based methods are almost unaffected by the choice of *k* (fig. 1). Unstable results are only observed for either small or high values of *k*, with respect to the total number of genomic regions analyzed. In comparison with established methods, such as pcadapt, the kNN-based methods do remarkably well. As expected, the *F*_*ST*_ results are fully comparable for star-like genealogies (fig. 1A). However, as soon as hierarchical structure is introduced to the population history, our proposed competing methods show overall higher AUC values. In fact, the kNN-based techniques remain almost unaffected when varying the coalescent times to the ancestral population *P*_*anc*_ (fig. 2A).

**FIG. 1.**
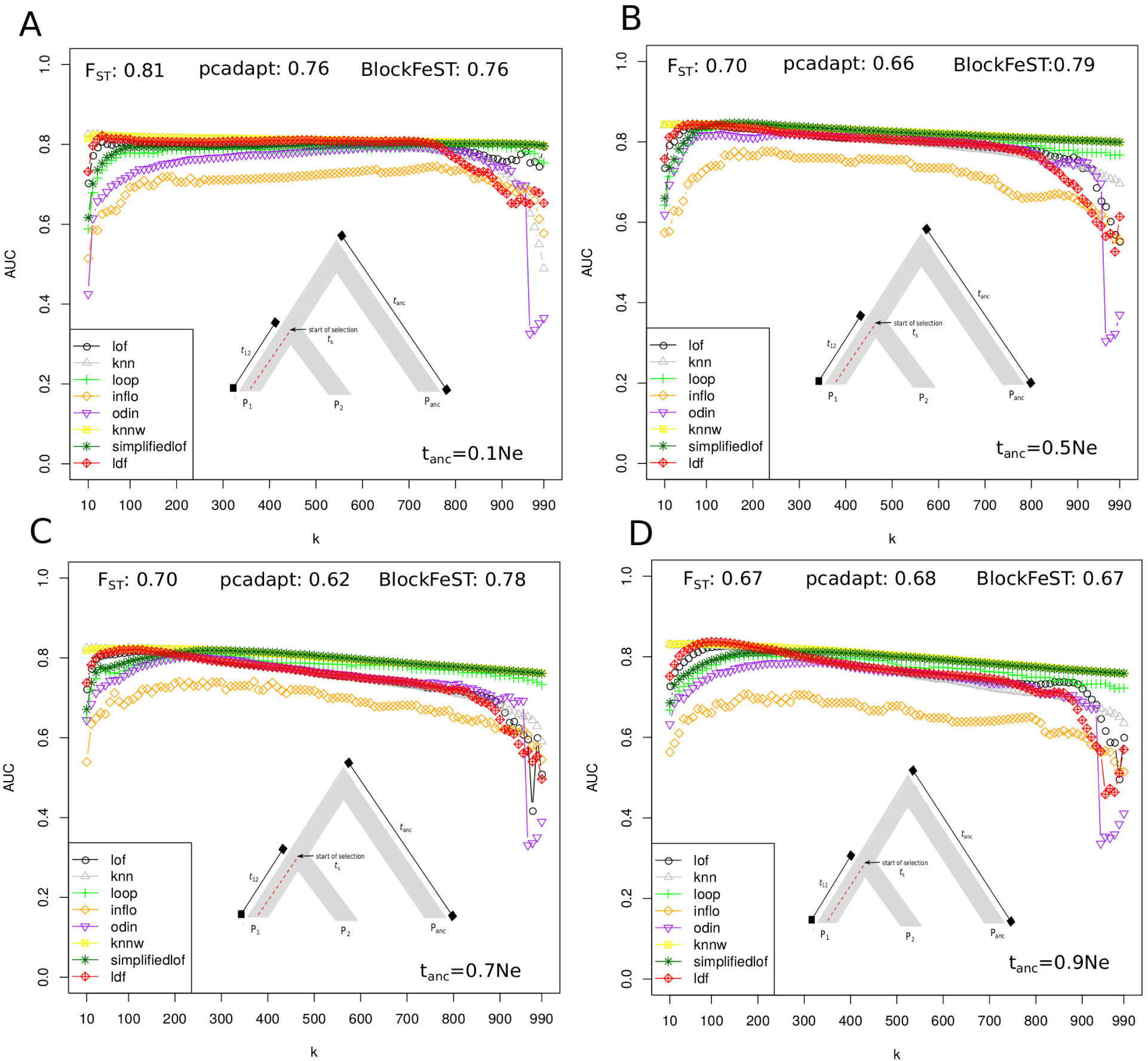
Positive directional selection: varying the coalescent time to the ancestral population (*t*_*anc*_). The results for the kNN-based methods using pairwise *F*_*ST*_ as features are shown for 100 sequentially sampled *k*s (*k*=[1,10*,…*,990,1000]) and in comparison to the accuracy of *F*_*ST*_, pcadapt and BlockFeST. Recombination rate is set to *r*=0.001. A. The simulations are based on a star formed genealogie (*t*_12_=0.1 = *t*_*anc*_). B. The coalescent time to the ancestral population is *t*_*anc*_=0.5*N*_*e*_. C. The coalescent time to the ancestral population is *t*_*anc*_=0.7*N*_*e*_. D. The coalescent time to the ancestral population is *t*_*anc*_=0.9*N*_*e*_.

**FIG. 2.**
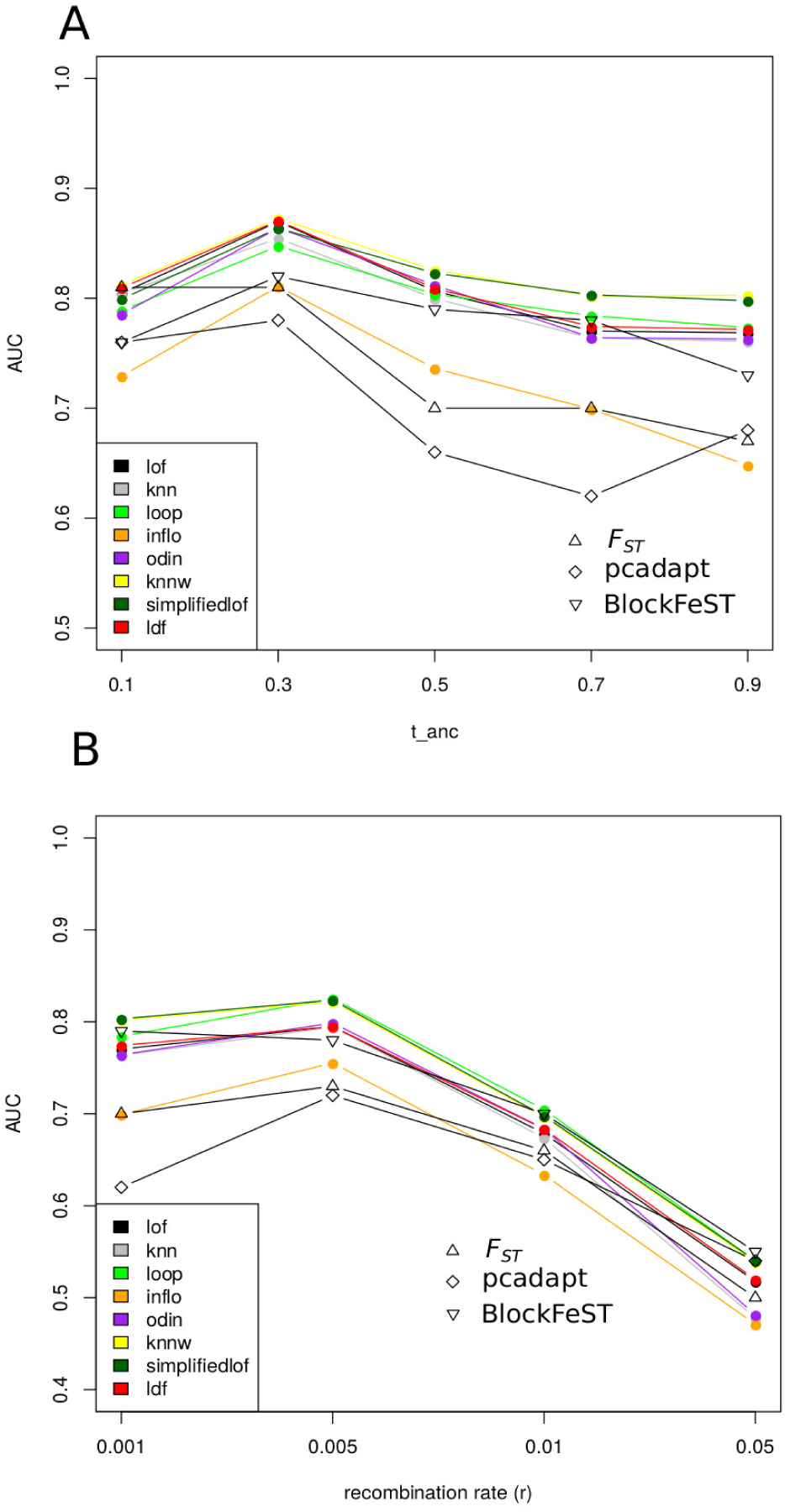
Detecting positive directional selection with a computed *k*. The kNN methods using pairwise *F*_*ST*_ as features compared to *F*_*ST*_, pcadapt and BlockFeST. A. Varying the coalescent time with the ancestral population (*t*_*anc*_=[0.1,0.3,0.5,0.7,0.9]*N*_*e*_ generations ago). The recombination rate is set to *r*=0.001. B. Varying the recombination rates (*r*=[0.001,0.005,0.01,0.05]). The coalescent time with the ancestral population is *t*_*anc*_=0.7*N*_*e*_ generations ago.

The weighted-kNN and simplified-LOF methods are the strongest kNN-based methods, both outperforming *F*_*ST*_ and pcadapt, and are comparable to BlockFeST (fig. 1, fig. 2). However, BlockFeST is based on computationally intensive MCMC runs and for that reason might not be generally applicable. Overall, the performance of all methods under consideration decreases with increasing recombination rates (fig. 4B). This is expected because the signal of selection gets eroded, which makes it harder to detect these patterns. Based on our simulations, we observed that the INFLO algorithm is the weakest kNN-based method for the detection of positive directional selection, especially when the recombination rates are high. Finally, INFLO is the most sensitive to background population histories as seen from fig. 2A.

### On the Power to Detect Introgression

Simulations under uni-directional introgression (see ‘Material and Methods’) from an archaic population (*P*_*anc*_) to an in-group population (*P*_2_) confirm that the kNN-based family of methods is almost unaffected by the choice of *k*. Surprisingly, we observe that in some cases *F*_*ST*_ outperforms the other more specialized methods (fig. 3A). However, we also report unstable results for *F*_*ST*_ when varying the time of gene-flow (fig. 4B). *D*_3_ is more stable in these cases and also outperforms the kNN-based methods based on pairwise *F*_*ST*_ features (fig. 4B).

**FIG. 3.**
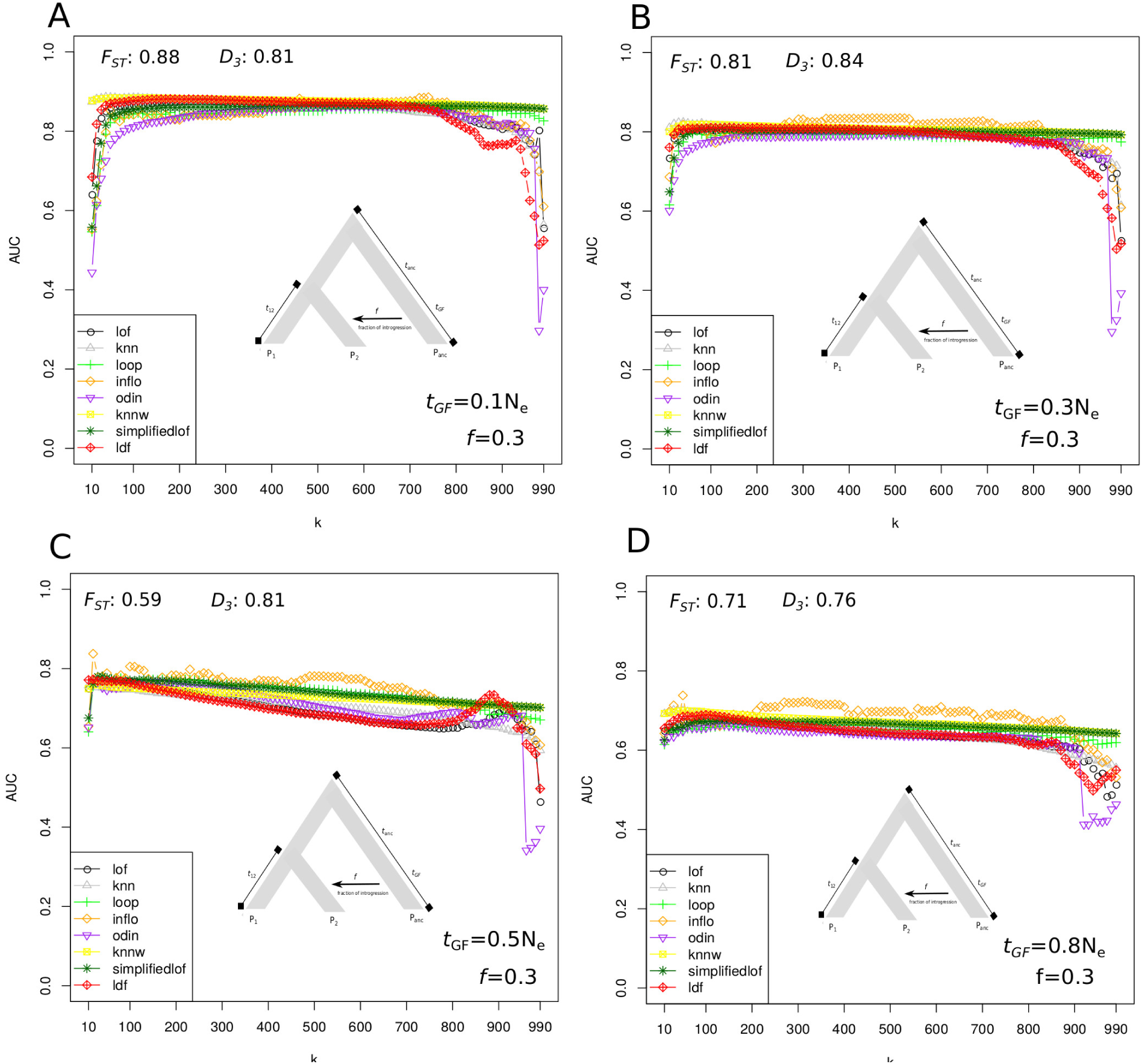
Varying the time of gene-flow (*t*_*GF*_). The results for the kNN-based methods using pairwise *F*_*ST*_ as features shown for 100 sequentially sampled *k*’s (*k*=[1,10*,...*,990,1000]). Coalescent times are *t*_12_=1*N*_*e*_ and *t*_*anc*_=2*N*_*e*_ generations ago. Recombination rate is set to *r*=0.01 in all simulations. The outcome of the kNN-based methods are compared to *F*_*ST*_ and *D*_3_. The time of gene-flow is set to A. *t*_*GF*_=0.1*N*_*e*_ B. *t*_*GF*_=0.3*N*_*e*_ C. *t*_*GF*_=0.5*N*_*e*_ and D. *t*_*GF*_=0.8*N*_*e*_ generations ago.

**FIG. 4.**
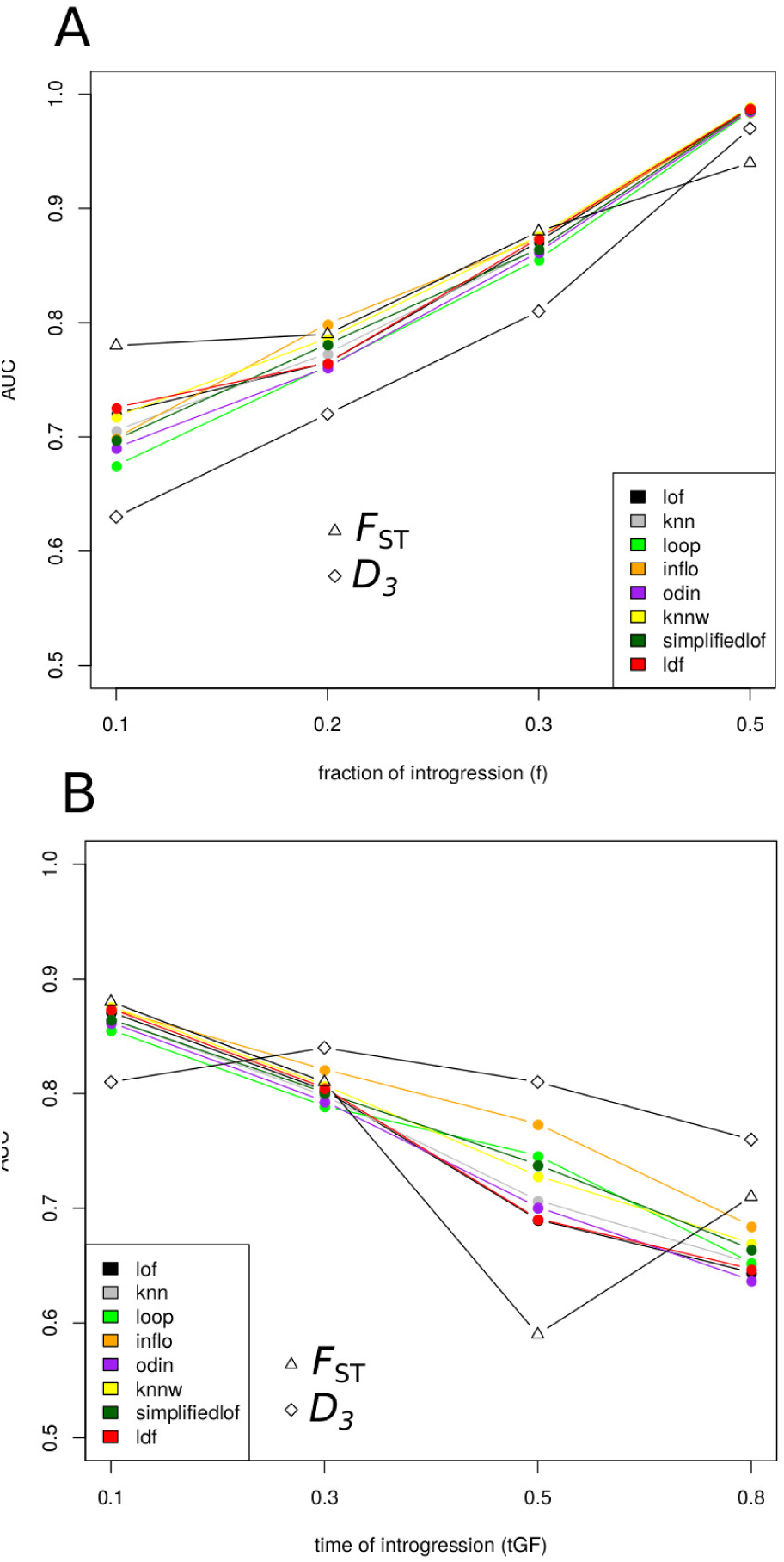
Detecting introgression with a computed *k*. The accuracy of the kNN-methods using pairwise *F*_*ST*_ as features compared to *F*_*ST*_ and *D*_3_. Recombination rate is set to *r*=0.01 in all simulations. A. Varying the fraction of introgression (*f*=[0.1,0.2,0.3,0.5]) B. Varying the time of gene-flow (*t*_*GF*_=[0.1,0.3,0.5,0.8]).

Increasing the proportion of introgression has the same effect on all kNN-based methods: the accuracy increases and is nearly at 100% when *f*=0.5. When gene-flow is recent *D*_3_ has overall lower AUC values than the kNN-based approaches (fig.4A). Interestingly, while INFLO is the weakest kNN-based method for the detection of directional selection, our simulations point at INFLO as a potentially powerful algorithm to detect local signatures of introgression (fig. 3). Overall, similar as reported for the positive directional selection cases, the weighted kNN method and simplified-LOF show high accuracy also in the introgression cases and provide stable score rankings almost across the full range of *k*. Also, in comparison to the other kNN approaches, low and very high *k* values have the most negative effect on ODIN and LDF.

Using *d*_*xy*_ as features also give stable results for almost all choices of *k* (Supplementary Fig. S1). However, in this situation results are not as good as those of kNN techniques with incorporated pairwise *F*_*ST*_ estimates. This is especially true when the time of gene-flow is recent, and a low fraction of introgression is shared by *P*_2_ and the archaic population *P*_*anc*_ (Supplementary Fig. S2).

### Application to the 1000 Genomes Data

We also analyzed the 1000 Genomes Data (1000 Genomes Project Consortium and others, 2015) to demonstrate the efficacy of our proposed kNN-based approaches when processing real data. The employed dataset is currently one of the largest publicly available datasets, both in terms of number of samples and number of SNPs, with 2,504 human samples from 26 populations, and 77,832,252 SNPs in the entire set of autosomes (phase 3). We applied all implemented kNN-based techniques on a per-autosome basis on the samples of the populations CEU, CHB, and YRI, evaluating non-overlapping sliding windows of size 100kb. Here, we summarize the results for chromosome 2 by reporting the windows all kNN tools agree on to be outlier windows. We report on the nearest genes to these outlier windows (Table 1). For each tool we consider kNN scores within a conservative 0.005-quantile to define the outlier candidates.

**Table 1.** Human chromosome 2 outlier windows

| Mb (start) | Mb (end) | Nearest genes | $F_{ST}$ | $\Delta F_{ST}$ |
| --- | --- | --- | --- | --- |
|  |  |  | [CEU/CHB,CEU/YRI,CHB/YRI] | [CEU/CHB,CEU/YRI,CHB/YRI] |
| 17.3 | 17.4 | VSNL1, AC010880.1 | [0.55,0.16,0.34] | [0.46,0.02,0.19] |
| **72.5** | **72.6** | **EXOC6B\*** | [0.15,0.45,0.69] | [0.06,0.31,0.54] |
| **72.6** | **72.7** | **EXOC6B\*** | [0.13,0.43,0.65] | [0.03,0.29,0.50] |
| 72.8 | 72.9 | EXOC6B* | [0.13,0.34,0.57] | [0.03,0.20,0.42] |
| 74.7 | 74.8 | CCDC142*, M1AP* | [0.53,0.37,0.22] | [0.43,0.23,0.07] |
| 104.1 | 104.2 | LINC01127 | [0.42,0.10,0.52] | [0.37,-0.04,0.36] |
| 109.1 | 109.2 | GCC2*,LIMS1* | [0.43,0.09,0.58] | [0.33,-0.05,0.43] |
| 109.2 | 109.3 | LIMS1* | [0.40,0.09,0.55] | [0.30,-0.04,0.40] |
| 109.5 | 109.6 | **EDAR\*** | [0.63,0.32,0.34] | [0.53,0.18,0.19] |
| 195.6 | 195.7 | LINC01790* | [0.03,0.50,0.42] | [-0.07,0.36,0.27] |

NOTE.—Shown are the 0.005-quantile outlier 100kb windows all kNN-based methods agree with and the nearest genes to these windows. The medoid $F_{ST}$ vector is $F_{ST}\text{-medoid} = [\text{CEU/CHB}=0.09, \text{CEU/YRI}=0.14, \text{CHB/YRI}=0.15]$ . The top-3 outlier windows are highlighted in bold. \*The outlier window overlaps with the gene

Describing the properties and attributes of all these genes may lead to the story-telling fallacy (Pavlidis *et al.*, 2012). We therefore report for some of them what has been reported in literature. The top-2 candidate genes for directional positive selection are the protein coding genes EXOC6B and EDAR (table 1, fig. 5). Baye *et al.* (2009) report EXOC6B as a positively selected gene. Intellectual disability and developmental delay are associated with this gene. Our kNN approaches suggest directional selection between the YRI population and both the CEU and CHB populations (table 1). Bryk *et al.* (2008) report EDAR, which is a gene involved in ectodermal development, increased in frequency in East Asia due to positive selection 10,000 years ago. The kNN-based approaches suggest the strongest effect between CEU and CHB (table 1).

**FIG. 5.**
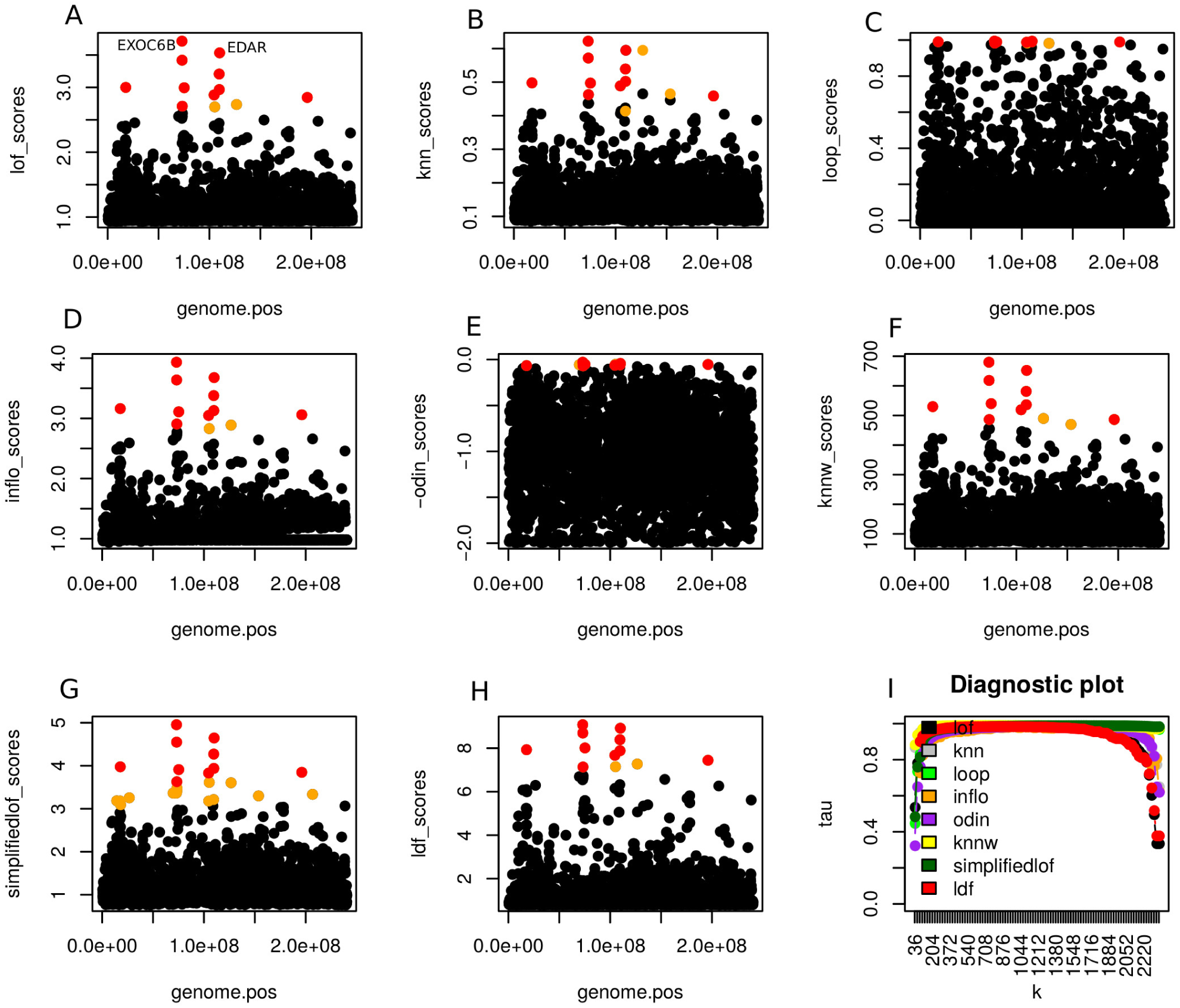
Genome scan plots of human chromosome 2. A-H. The kNN scores are shown along human chromosome 2 based on 100kb consecutive sliding windows. Red and Orange dots are the outliers found by that specific kNN tool (0.005-quantile of the scores). Red dots indicate that all kNN tools agree on these outliers. I. A diagnostic plot is shown with the pairwise rank correlations of the kNN scores while varying the parameter *k*.

Another candidate gene is CNTNAP5 (outlier window: 126.1-126.2Mb) and is confirmed by all tools but ODIN. The selection effect is ∆*F*_*ST*_ = [CEU/CHB=0.39, CEU/YRI=0.01, CHB/YRI=0.25] suggesting directional selection in the Asian population (CHB). An additional candidate gene is FMNL2 (outlier window: 153.1-153.2Mb) and is exclusively reported by the weighted-kNN and kNN algorithms with a selection effect of ∆*F*_*ST*_ =[CEU/CHB=0.29, CEU/YRI=0.11, CHB/YRI=0.33]). The genomic region 104.7-104.8Mb is reported by all tools but the weighted-kNN and kNN. The nearest gene is LINC01127 and the selection effect is ∆*F*_*ST*_ =[CEU/CHB=0.32, CEU/YRI=0.21, CHB/YRI=-0.10]. Finally, the ODIN method exclusively reports on the ANTXR1 gene (outlier window: 69.2-69.3Mb) as a candidate for selection with a selection effect of ∆*F*_*ST*_ =[CEU/CHB=0.22, CEU/YRI=0.31, CHB/YRI=-0.05] slightly pointing to positive directional selection in the european population (CEU) and a reduced diversity between CHB and YRI.

## Discussion

In this paper, we have investigated the usage of the kNN-based algorithms to detect local signatures of selection and introgression in whole-genome scans. Coalescent simulations under positive directional selection and introgression show that the kNN-based methods using *F*_*ST*_ as features perform remarkably well, and are almost unaffected by the choice of *k*. Also, the approaches presented in this work are highly flexible with regard to the choice of the feature set and researcher are not limited to using *F*_*ST*_ or *d*_*xy*_. A different set of features, or a combination of selection/introgression sensitive features may further improve the accuracy. Furthermore, in contrast to other genome-scan approaches, the kNN-based approaches are based on simple concepts while at the same time do not depend on specific assumptions about the distributions of the underlying data. The algorithm implemented in the R-package pcadapt, for example, uses a principal component transformation of the data in combination with a linear regression model, and thus assumes linear relationships between populations.

We have demonstrated that the evaluated kNN-based methods achieve qualitatively comparable performance with the Bayesian approach implemented in the R-package BlockFeST when detecting positive directional selection, while being considerably less compute-intensive. We showcased the capacity of the kNN-based methods to analyze real-world data by scanning the second chromosome of the human genome (data available by the 1000 Genomes project). We confirm known genes under positive selection, like EDAR and EXOC6B, but also report a set of new candidate genes, like LIMS1 and CNTNAP5. Outlier loci with significantly reduced diversity, and thus potentially pointing to gene-flow or balancing selection cannot be reported for human chromosome 2. The only candidate genes showing that type of signal are the LINC01127 and ANTXR1 genes, with slightly reduced diversity between the CHB and YRI populations.

We have also discussed certain challenges that arise when employing kNN-based techniques. A widely known complication with the kNN-based methods is the choice of *k*, for which the optimal value highly depends on the data. This problem is not fully solved with our approaches. We have shown, however, that under coalescent simulations and a convincing set of population models, the parameter *k* does not greatly affect the accuracy of our approaches. Future investigations will analyze the power of the kNN techniques, both analytically as well as through additional simulations over a wide range of population models and feature sets.

## Materials and Methods

### Simulation of Positive Directional Selection

We generated 950 neutral regions and 50 regions under positive directional selection (fig. 6) with the MSMS software tool (Ewing and Hermisson, 2010). The main calls to the MSMS program are as follows

Neutral model:

**FIG. 6.**
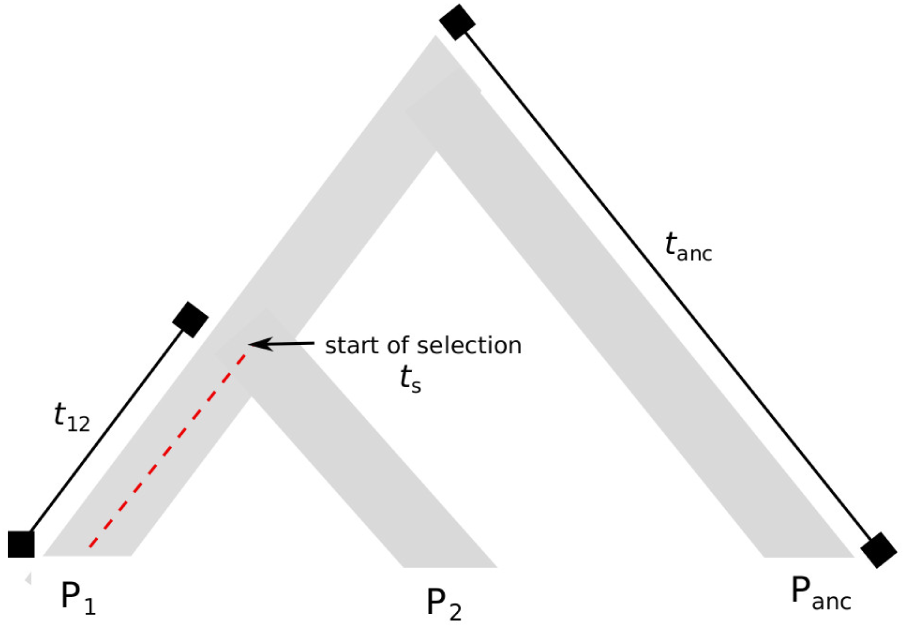
A sketched graphical illustration of positive directional selection. A three population genealogy with positive directional selection introduced at *t*_*s*_*N*_*e*_ generations ago in population *P*_1_.

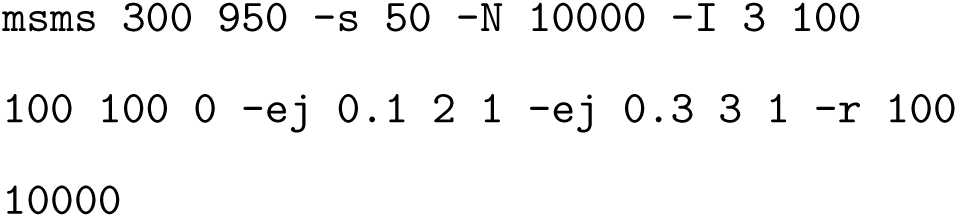

Alternative model:

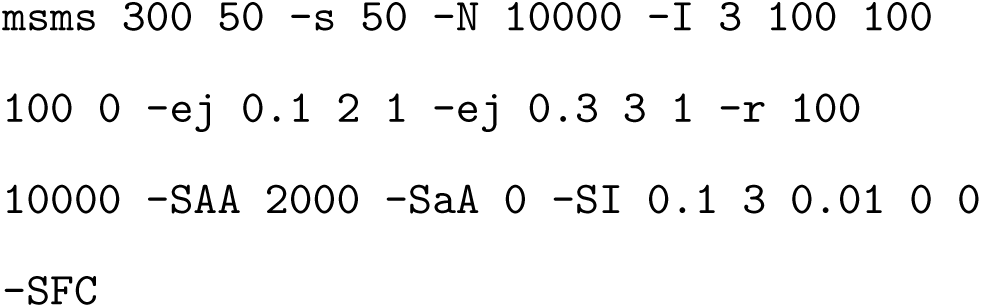

The above calls generate three populations, each comprising 100 samples (`-I`). The number of SNPs per each region is 50 (`-s`), and the effective population size is *N*_*e*_ =10000 (`-N`). The first coalescent event of population *P*_1_ and *P*_2_ is fixed at *t*_12_=0.1*N*_*e*_, and the second coalescent event is set to *t*_13_=0.2*N*_*e*_=*t*_*anc*_ generations ago. The selection strength for homozygotes is *s*=0.1 (`-SAA`), where selection starts at *t*_*s*_=0.1*N*_*e*_ generations ago (`-SI`) in population *P*_1_. The recombination rate is *r*=0.01 (-r). We varied recombination rates (*r*=[0,0.001,0.005,0.01,0.05]) and the time of coalescence with the ancestral population (*t*_*anc*_=[0.1,0.3,0.5,0.7,0.9]). In each of these simulations, we made use of the SFC parameter in order to drop simulations when the selected allele gets lost.

### Simulation of Introgression

To generate introgression events (fig. 7) we follow the simulation set-up of Martin *et al.* (2014). The below calls to the MS software (Hudson, 2002) generate the topologies:

**FIG. 7.**
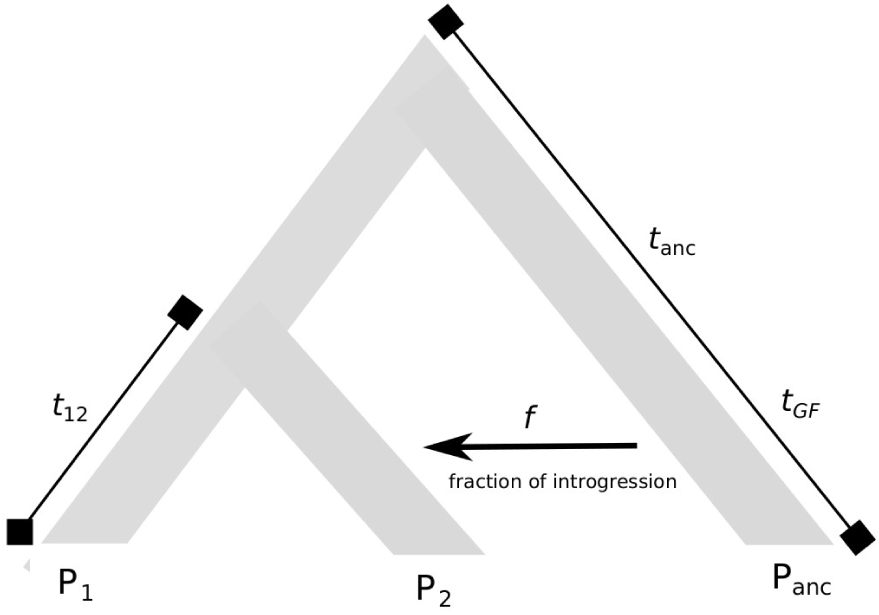
A sketched graphical illustration of introgression. A three population species tree with an uni-directional introgression event from the ancestral population *P*_*anc*_ to population *P*_2_ introduced *t_GF_ N*_*e*_ generations ago. The proportion of introgression is indicated by *f*.

Neutral model:

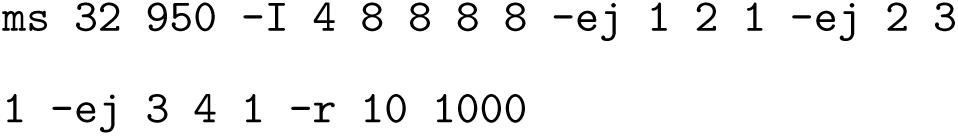

Alternative model:

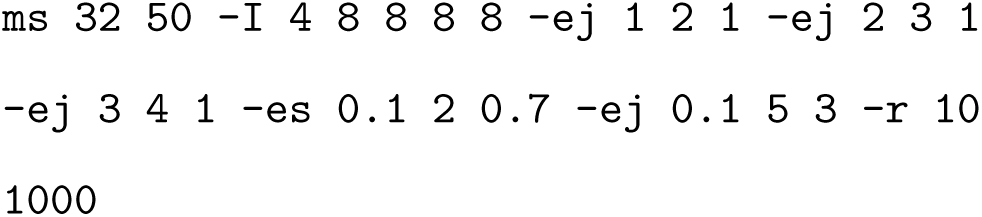

These calls simulate 950 neutral regions and 50 regions under introgression with four populations including 8 samples each (`-I`). The coalescent times are *t*_12_=1*N*_*e*_, *t*_13_=2*N*_*e*_=*t*_*anc*_ and *t*_14_=3*N*_*e*_=*t*_*O*_ generations ago. We introduced *P_anc_ → P*_2_ introgression *t*_*GF*_=0.1*N*_*e*_ generations ago (`-es`) with a fraction of introgression *f*=0.3 (`-es 0.1 2 [1-f])`. The recombination rate was set to *r*=0.01 in all simulations.

Finally, the nucleotide sequences were generated using the seq-gen (Rambaut and Grass, 1997) software with the following call:

~~~
seq-gen -mHKY -I 1000 -s 0.01
~~~

This generates 1kB sequences under the Hasegawa-Kishino-Yano (`-mHKY`) substitution model with a branch scaling factor of *s*=0.01 (`-s`). We varied the proportion of introgression (*f*=[0.1,0.2,0.3,0.5]) and the time of gene-flow (*t*_*GF*_=[0.1,0.3,0.5,0.8]).

### Comparison with Other Methods

We selected a set of kNN-based techniques similar to Campos *et al.* (2016) and as implemented in the ELKI software (Schubert and Zimek, 2019). For the selection cases, we contrast the kNN-based algorithms while incorporating pairwise *F*_*ST*_ as features to the method implemented in the R-package pcadapt (Luu *et al.*, 2017). We computed the sum of *log-p-values* to label a region to make it comparable to the other methods under consideration. The number of principal components was set to K=2. In addition, we report the accuracy of the recently published method implemented in the R-package BlockFeST (Pfeifer and Lercher, 2018) using the default parameters.

In the introgression cases we compare the kNN-based methods to the *D*_3_ approach (Hahn and Hibbins, 2019) which follows a similar logic as the *d*_*f*_ method introduced by Pfeifer and Kapan (2019), but without the need of an outgroup. This method is compared to the kNN-based techniques when using pairwise *d*_*xy*_ and pairwise *F*_*ST*_ as features. Finally, we relate all of these methods to the *F*_*ST*_ estimate (Hudson *et al.*, 1992) as a baseline approach. Accuracy is measured by the Area Under the Curve (AUC), as implemented in the R-package pROC (Robin *et al.*, 2011).

### Code Availability

We provide R scripts to perform kNN-based whole genome scans, available at the GitHub repository *pievos101/kNN-Genome-Scans*. The code interfaces with the powerful genomics R-package PopGenome (Pfeifer *et al.*, 2014), and enables flexible genomic scans with sliding windows as well as genomic scans based on genomic features such as genes, UTRs or exons.

## Supporting information

Supplement Figures Revised

## Acknowledgments

We thank Fernando Racimo and Ben Rosenzweig for their valuable comments on our manuscript.

## Notes

#### Summary of Updates

RNDmin was not correctly included... (Thanks Ben!)

