## Supplement Figures Revised for "Genome Scans for Selection and Introgression based on *k*-nearest Neighbor Techniques"

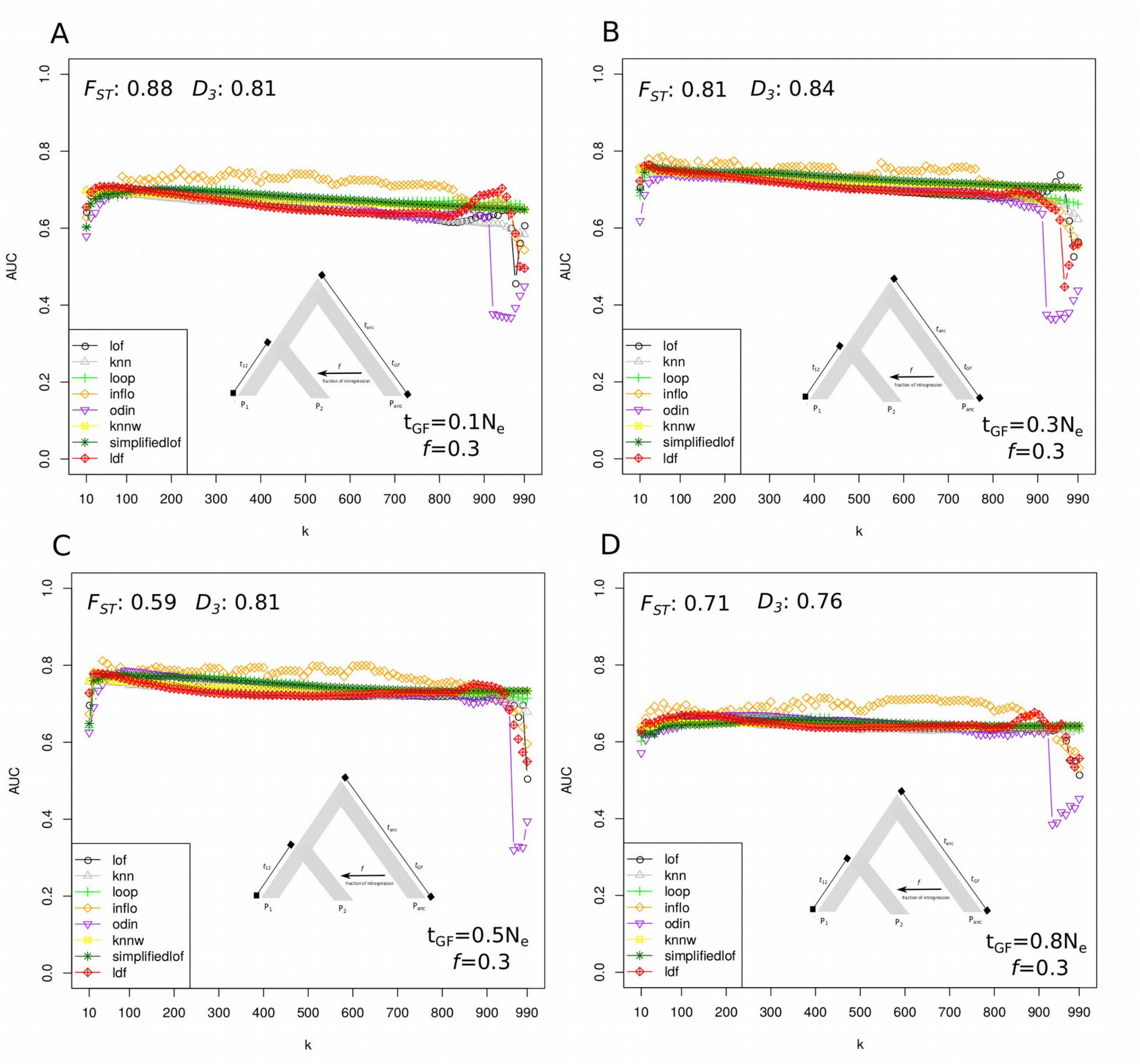


**Supplementary Figure S1: Varying the time of gene-flow (*$t_{GF}$*) and using *$d_{xy}$* as features.** The results for the *kNN*-based methods using $d_{xy}$ as features shown for 100 sequentially sampled *k's* (*k*=[1, 10, … ,990, 1000]). The coalescent times are $t_{12}$=1$N_e$ and $t_{anc}$=2$N_e$ generations ago. Recombination rate is set to *r*=0.01 in all simulations. The outcome of the *kNN*-based methods are compared to $F_{ST}$ and $D_3$. The time of gene-flow is set to **A.** $t_{GF}$=0.1$N_e$ **B.** $t_{GF}$=0.3$N_e$ **C.** $t_{GF}$=0.5$N_e$ and **D.** $t_{GF}$=0.8$N_e$ generations ago.

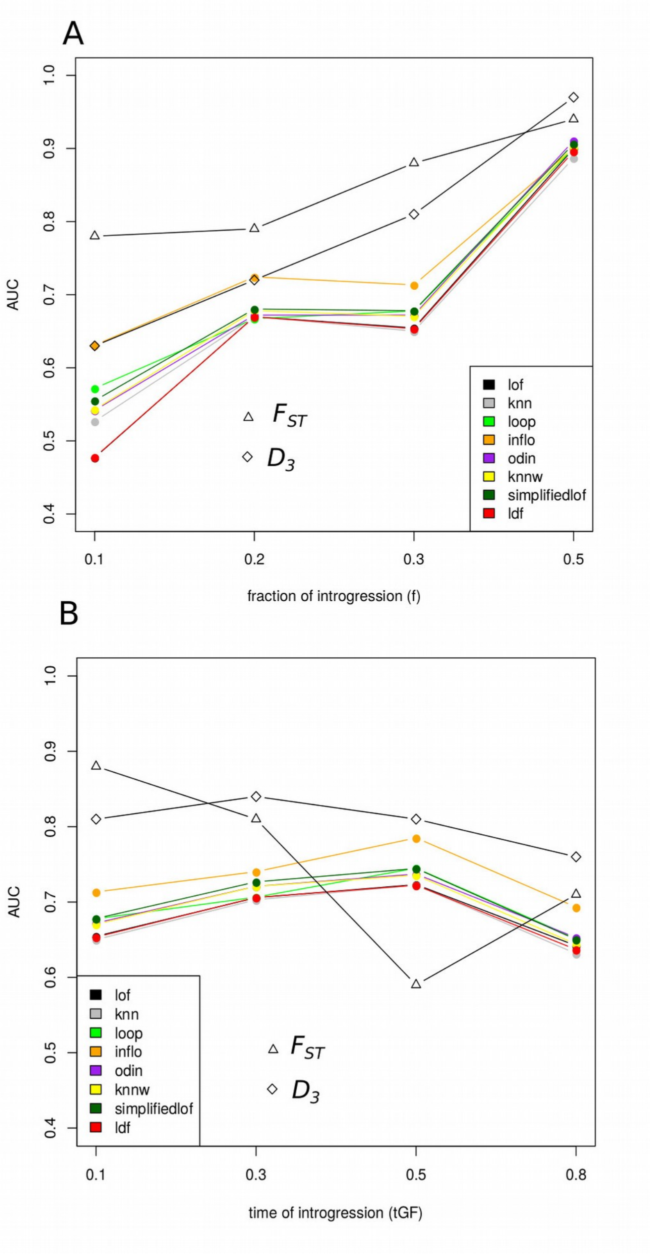


**Supplementary Figure S2: Detecting introgression with a computed *k* and using $d_{xy}$ as features.** The accuracy of the *kNN*-methods using $d_{xy}$ as features compared to $F_{ST}$ and $D_3$. Recombination rate is set to $r$=0.01 in all simulations. **A.** Varying the fraction of introgression ($f$=[0.1, 0.2, 0.3, 0.5]) **B.** Varying the time of gene-flow ($t_{GF}$=[0.1, 0.3, 0.5, 0.8]).
